# Locating suspicious lethal genes by abnormal distributions of SNP patterns

**DOI:** 10.1101/530733

**Authors:** Xiaojun Ding

## Abstract

A gene, a locatable region of genomic sequence, is the basic functional unit of heredity. Differences in genes lead to the various congenital physical conditions of people. One kind of these major differences are caused by genetic variations named single nucleotide polymorphisms(SNPs). SNPs may affect splice sites, protein structures and so on, and then cause gene abnormities. Some abnormities will lead to fatal diseases. People with these diseases have a small probability of having children. Thus the distributions of SNP patterns on these sites will be different with distributions on other sites. Based on this idea, we present a novel statistical method to detect the abnormal distributions of SNP patterns and then to locate the suspicious lethal genes. We did the test on HapMap data and found 74 suspicious SNPs. Among them, 10 SNPs can map reviewed genes in NCBI database. 5 genes out of them relate to fatal children diseases or embryonic development, 1 gene can cause spermatogenic failure, the other 4 genes are also associated with many genetic diseases. The results validate our idea. The method is very simple and is guaranteed by a statistical test. It is a cheap way to discover the suspicious pathogenic genes and the mutation site. The mined genes deserve further study.

**Author summary:** Xiaojun Ding received the BS, MS and PhD degrees in computer science from Central South University. Now he is a assistant professor in Yulin Normal University. His research interests include computational biology and machine learning.

## Introduction

Genes are the most important genetic materials which can determine the health of a person in some ways. The functions of genes may be affected by the genetic variations called SNPs. So it is a good way to study the disease-related genes from SNPs. Many defective genes caused by SNPs for human Mendelian diseases (i.e. single gene diseases) have bAsnaghi2013Ieen found [1, 2]. For example, Prescott el al. [3] found a nonsynonymous SNP in ATG16L1 related to Crohn’s disease. Seki et al. [4] reported that a functional SNP in CILP is suspicious to lumbar disc disease. These achievements inspire people.

However, the discovered pathogenic genes caused by SNPs only take a small fraction, most of them are still unknown. At the same time, the number of SNPs is very large and most SNPs do not take effects on genes [5, 6]. Checking all SNPs by biological experiments is a expensive work. Narrowing the range of suspicious SNPs will benefit the study of pathogenic genes greatly [7]. For the purpose, people analyzed SNPs from various angles. Lee et al. [8] builded a functional SNP database which integrates information got from 16 bioinformatics tools and functional SNP effects for disease researches. Cargill et al. [9] studied the different rates of polymorphism within genes and between genes. They concluded that the rates may reflect selection acting against deleterious alleles during evolution and the lower allele frequency of missense cSNPs are possibly associated with diseases. Adzhubei et al. [10] developed a tool named PolyPhen which predicts possible impact of an amino acid substitution on the structure and function of a human protein. Kumar et al. [11] developed a tool named SIFT which predicts whether an amino acid substitution will affect protein function. Their algorithm is suitable to naturally occurring nonsynonymous polymorphisms and laboratory-induced missense mutations. While synonymous mutations can also contribute to human diseases [12]. For example, Westerveld et al. [13] reported that a intronic variants rs1552726 may affect the splice site activity.

In the paper, a novel method is proposed from the angle of genetic law. If a defective gene caused by SNPs can lead to fatal diseases and most of the sick people do not have the change to breed the next generation. This will affect the distributions of the SNPs within the gene. It provides us a novel way to distinguish the pathogenic SNPs from normal SNPs.

## Materials and methods

Given a bi-allele SNP, ‘A’ and ‘a’ are used to denote the major and minor allele, respectively. Because chromosomes come in pairs, each individual will take one of the following three SNP patterns: pattern0=‘AA’, pattern1=‘Aa’, pattern2=‘aa’. In a population, the distribution of individuals taking each pattern can be counted. The abnormal distributions are what we concern. Next, an example is given to illustrate the abnormal distributions.

Example 1: Suppose that there is a distribution for 1000 individuals; 500 individuals out of them take pattern ‘AA’ and other people take pattern ‘aa’. Nobody takes the pattern ‘Aa’.

According to bisexual reproduction rule, a child will inherit one chromosome from his mother and one from his father. If the mother takes pattern ‘AA‘ and the father takes pattern ‘aa‘. The child will take pattern ‘Aa‘. It is shown as Fig 1. If every one has a equal probability to marry other people in the population. The probability of a child with pattern ‘Aa’ should be 2*0.5*0.5=0.5, that means there should be about 0.5*1000=500 individuals taking the pattern ‘Aa‘. But none is observed. We think that the distribution on this SNP site is abnormal. The reason for the abnormal distribution is probably that the person taking ‘Aa’ will die in childhood so that we can not observe them. From the analysis, a hypothesis is proposed as following.

**Fig 1.**
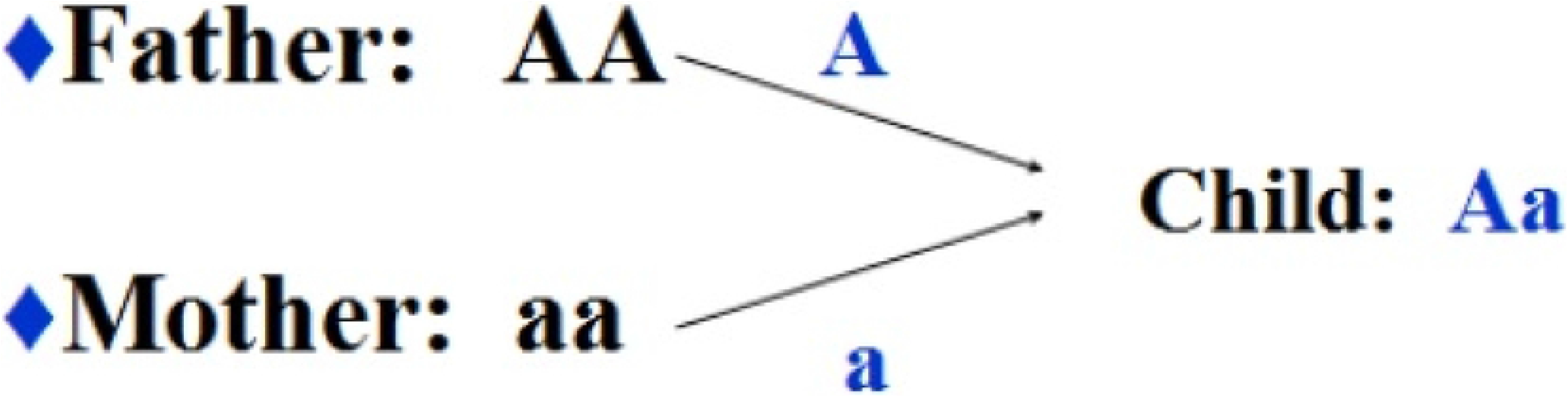
The heredity of SNPs.

### Hypothesis 1

Some SNP patterns should appear in human populations according to bisexual reproduction but are not observed. These patterns may cause gene abnormities for fatal genetic diseases so that the corresponding person will die in childhood.

In HapMap data, the SNP data of 11 human populations are sequenced. Since the relationships of individuals in each population are unknown, we make a assumption to simplify the computation.

### Assumption 1

In each population, everyone has the equal probability to marry other people and to give a birth to a baby.

For *population_j_*, the distribution *P* of individuals for all the patterns can be counted. *P* = [*p*_0_,*p*_1_,*p*_2_], where *p_i_* is the percentage of the individuals with *pattern_i_*.Under Assumption 1 and the bisexual reproduction rule, the distribution among the next generation (denoted by *P*\*) can be computed according to the distribution *P*. Let 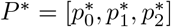. *P\** can be computed by the formulas (1),(2) and (3).

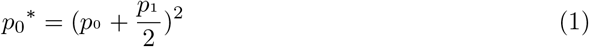

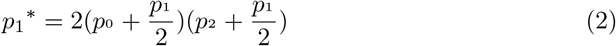

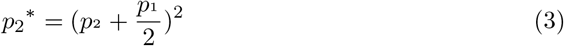

If there is no big disaster, the distribution among a human population will not change radically. Under usual circumstances, *P*\* can be treated as an approximation to the mean distribution of the current population. If *p_i_* is 0, but 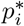 is far from 0. The distribution may be abnormal. Supposing the size of the *population_j_* is *n_j_*. The number of individuals matching the *pattern_i_* obeys the binomial distribution. *e_ij_* is used to denote the event that the *pattern_i_* is not observed in current *population_j_*, the probability of *e_ij_* is computed by formula (4)

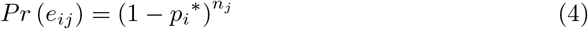

In HapMap data, there are 11 human populations. *eAll* denotes the event that the *pattern_i_* can not be observed in all the populations. The probability of *eAll* is given by formula (5).

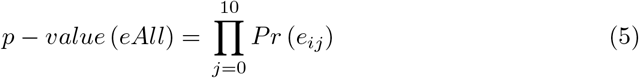

If the *p* – *value(eAll*) is very small, the event that the *pattern_i_* are not observed in all the populations is unlikely. But it actually happens in the observation of HapMap project. The reason may be that the *pattern_i_* may cause gene abnormities for fatal genetic diseases and most patients will die in childhood. In usual statistical test, 0.05 is often used as the threshold of significance level. In our test, since there are many SNPs to be checked. For example, there are 117068 SNPs on chromosome 1. The significance level should be corrected. Here, Bonferroni correction is choosed to adjust the threshold. Given *k* SNPs, 3*k* hypothesizes need to be tested. The p-value threshold of significance should be 0.05/3*k*. The SNPs with *p* – *value(eAll*) below the p-value threshold are suspicious. After finding these SNPs, the suspicious lethal genes can be located by NCBI database.

The HapMap [14] data (genotypes data of phase 3.3 consensus) is used to validate our method. The HapMap data contains 11 human populations. 1417 individuals are sequenced. The detail of human populations is listed in Table 1.

**Table 1.** information of human populations.

| Label | Population sample | Size |
| --- | --- | --- |
| ASW | African ancestry in Southwest USA | 87 |
| CEU | Utah residents with Northern and Western European ancestry from the CEPH collection | 174 |
| CHB | Han Chinese in Beijing, China | 139 |
| CHD | Chinese in Metropolitan Denver, Colorado | 109 |
| GIH | Gujarati Indians in Houston, Texas | 101 |
| JPT | Japanese in Tokyo, Japan | 116 |
| LWK | Luhya in Webuye, Kenya | 110 |
| MEX | Mexican ancestry in Los Angeles, California | 86 |
| MKK | Maasai in Kinyawa, Kenya | 184 |
| TSI | Toscans in Italy | 102 |
| YRI | Yoruba in Ibadan, Nigeria | 209 |

## Results

In our experiment,the statistical test is checked on each chromosome separately. For one chromosome, the common bi-allele SNPs in the 11 populations are extracted. On chromosome 1, there are 117068 common SNPs and 8 SNPs are detected to be suspicious. On the whole 22 chromosomes, 74 SNPs are obtained among 1,395,560 SNPs (we do not check SNPs on the chromosome X and Y because the heredity on chromosome X and Y are different with that on autosomes). By looking up in NCBI database, 29 genes are located. Some of them have been studied in many years and some are still new to people. According to the reference sequence (RefSeq) status code, these genes can be divided into 3 catalogs.

A reviewed gene means that its RefSeq record has been reviewed by NCBI staff or by a collaborator. The NCBI review process includes assessing available sequence data and the literature. Some RefSeq records may incorporate expanded sequence and annotation information.

In our results, 10 SNPs map reviewed genes. The corresponding chromosomes, SNPs, genes, gene types, alleles, disease patterns and p-values are listed in Table 2. For the first two SNPs and their patterns, the expectations of the number of individuals in each population are listed in Table 3 and Table 4, respectively. For the pattern ‘AA’ at SNP site rs2145402, there should be about 34.4 individuals in all populations. But none is observed. If the pattern ‘AA’ is normal, the probability of the event is only 3.05E-16. It is too small to happen by chance. So we think the distribution of the pattern at SNP rs2145402 is abnormal. The corresponding gene LYST is suspicious to genetic diseases.

**Table 2.** information of SNPs mapping reviewed genes.

| chr | SNP | gene | alleles | disease pattern | pvalue |
| --- | --- | --- | --- | --- | --- |
| 1 | rs2145402 | LYST | A/C | AA | 3.05E-16 |
| 1 | rs4915931 | ROR1 | A/G | AA | 1.70E-12 |
| 1 | rs4660992 | BMP8B | C/T | TT | 9.91E-10 |
| 6 | rs9263745 | CCHCR1 | A/G | AA | 5.67E-11 |
| 7 | rs11766679 | DPP6 | A/G | GG | 4.29E-10 |
| 10 | rs12263497 | INPP5F | A/G | GG | 1.46E-10 |
| 11 | rs1552726 | NLRP14 | A/G | GG | 2.64E-09 |
| 14 | rs3742943 | JAG2 | C/T | TT | 6.07E-08 |
| 16 | rs1646233 | CBFA2T3 | A/G | AA | 1.20E-09 |
| 22 | rs11705619 | TXNRD2 | C/T | CC | 1.43E-09 |

**Table 3.** expectations of the number of individuals in each population who takes pattern ‘AA‘ at SNP site rs2145402.

| population | CHB | CHD | JPT | CEU | TSI | ASW | LWK | MKK | YRI | GIH | MEX | All |
| --- | --- | --- | --- | --- | --- | --- | --- | --- | --- | --- | --- | --- |
| expectation | 1.1 | 0.4 | 0.5 | 11.6 | 8.2 | 1.0 | 0.0 | 0.1 | 0.1 | 4.8 | 6.6 | 34.4 |

**Table 4.** expectations of the number of individuals in each population who takes pattern ‘AA‘ at SNP site rs4915931.

| population | CHB | CHD | JPT | CEU | TSI | ASW | LWK | MKK | YRI | GIH | MEX | All |
| --- | --- | --- | --- | --- | --- | --- | --- | --- | --- | --- | --- | --- |
| expectation | 0.2 | 0.1 | 0.2 | 8.8 | 4.2 | 2.5 | 1.3 | 0.3 | 2.5 | 4.1 | 2.5 | 26.7 |

The reviewed genes are familiar to people. There are relatively more studies on them. The number of literatures focused on this kind of genes are more than that of other genes. These related literatures provide us a intuitive impression and are listed in the following. From the details we can see that most of these genes are actually associated with fatal genetic diseases.

1. SNP rs2145402 maps gene LYST. In ClinVar database [15], LYST is associated with lung cancer, Malignant melanoma and Chediak-Higashi syndrome. Many researchers [16–21] also reported that gene LYST is associated with Chediak-Higashi syndrome. Chediak-Higashi syndrome can affect many parts of the body, particularly the immune system. The disease damages immune system cells. Most affected individuals have repeated and persistent infections starting in infancy or early childhood [22]. The result of the disease is very serious and most affected individuals die in childhood [23].
2. SNP rs4915931 maps gene ROR1 In ClinVar database [15], ROR1 is associated with malignant melanoma. Broome et al. [24] reported that ROR1 is a receptor tyrosine kinase expressed during embryogenesis, on chronic lymphocytic leukemia and in other malignancies. Hudecek et al. [25] found that ROR1 is highly expressed during early embryonic development but expressed at very low levels in adult tissues. Many papers reported that ROR1 has a very close relation with chronic lymphocytic leukemia [26, 27] and acute lymphoblastic leukemia [28–32]. ROR1 is suggested as the targeted therapy for human malignancies [33, 34].
3. SNP rs4660992 maps gene BMP8B BMP8B is a thermogenic protein which increases brown adipose tissue thermogenesis through both central and peripheral actions and regulates energy balance in partnership with hypothalamic AMPK [35]. Zhao et al. [36] showed that mouse Bmp8a (Op2) and Bmp8b play a role in spermatogenesis and placental development. Ying et al. [37] reported that BMP8B is required for the generation of primordial germ cells in the mouse.
4. SNP rs11766679 maps gene DPP6 Genetic variation in DPP6 is associated with amyotrophic lateral sclerosis [20, 38, 39] and Familial Idiopathic Ventricular Fibrillation [40]. Golz et al. [41] found that DPP6 is suspicious to many diseases such as cardiovascular diseases, endocrinological diseases, metabolic diseases, gastroenterological diseases, cancer, hematological diseases, inflammation, muscle skeleton diseases, neurological diseases, urological diseases, reproduction disorders and respiratory diseases.
5. SNP rs12263497 maps gene Inpp5f Zhu el al. [42] reported that Inpp5f is a polyphosphoinositide phosphatase that regulates cardiac hypertrophic responsiveness. Kim et al. [43] found that INPP5F inhibits STAT3 activity and suppresses gliomas tumorigenicity. Palermo et al. [44] reported that gene expression of INPP5F can be as an independent prognostic marker in fludarabine-based therapy of chronic lymphocytic leukemia. Bai et al. [45] reported that alteration of Akt signal plays an important role in diabetic cardiomyopathy. Inpp5f is a negative regulator of Akt signaling.
6. SNP rs9263745 maps gene CCHCR1 CCHCR1 is associated with malignant melanoma in ClinVar database [15]. CCHCR1 is up-regulated in skin cancer and associated with EGFR expression [46]. The CCHCR1 (HCR) gene is relevant for skin steroidogenesis and downregulated in cultured psoriatic keratinocytes [47].
7. SNP rs1552726 maps gene NLRP14 NLRP14 may play a regulatory role in the innate immune system [48]. Mutations in the testis-specific NALP14 gene in men suffering from spermatogenic failure in GeneCards database [49]. Westerveld et al. [13] collected the data of 157 patients, they identified 25 suspicious variants in total: 1 nonsense mutation, 14 missense mutations, 6 silent mutations and 4 intronic variants. By using ESEfinder and SpliceSiteFinder to check these SNPs, only the SNP rs1552726 is predicted to affect the correct splicing. Abe et al. [50] reported that germ-cell-specific inflammasome component NLRP14 negatively regulates cytosolic nucleic acid sensing to promote fertilization.
8. SNP rs3742943 maps gene JAG2 GO annotations related to JAG2 include Notch binding and calcium ion binding. The gene serve as a ligand for Notch signaling receptors. The Notch signaling pathway is an intercellular signaling mechanism that is essential for proper embryonic development. Defect in JAG2 may cause ossifying fibroma and shipyard eye in GeneCards database [49]. Houde et al. [51] observed the overexpression of the NOTCH ligand JAG2 in malignant plasma cells from multiple myeloma patients and cell lines. Yustein et al. [52] validated that induction of ectopic Myc target gene JAG2 augments hypoxic growth and tumorigenesis in a human B-cell model. Asnaghi et al. [53] reported that a role for Jag2 promotes uveal melanoma dissemination and growth. Vaish et al. [54] reported that JAG2 enhances tumorigenicity and chemoresistance of colorectal cancer cells.
9. SNP rs1646233 maps gene CBFA2T3 CBFA2T3-GLIS2 Fusion Protein Defines an Aggressive Subtype of Pediatric Acute Megakaryoblastic Leukemia [55] and CBFA2T3-GLIS2 fusion transcript is a novel common feature in pediatric, cytogenetically normal AML, not restricted to FAB M7 subtype [56]. CBFA2T3-GLIS2-positive is closed related to pediatric acute megakaryoblastic leukemia with non-Down syndrome [57].
10. SNP rs11705619 maps gene TXNRD2 Mutations in the gene TXNRD2 cause dilated cardiomyopathy [58] Jakupoglu et al. [59] did the experiment of Txnrd2 deletion and found which leads to fatal dilated cardiomyopathy and morphological abnormalities of cardiomyocytes. Prasad et al. [60] reported that TXNRD2 knockout is embryonic lethal in mice due to cardiac malformation.

## Conclusion

In the paper, we gain enlightenment from Mendel’s genetic experiments and propose a simple method which can utilize the distributions of SNP patterns among human populations to mine the pathogenic genes. On HapMap data, 74 SNPs are selected in 22 autosomal chromosomes, 10 SNPs can map reviewed genes in NCBI database. Among these genes, LYST gene and ROR1 gene are reported to relate to fatal genetic children diseases. Genes JAG2, TXNRD2 and BMP8B play important roles in embryonic development and lead to many fatal diseases. NALP14 gene may cause spermatogenic failure. Among 25 suspicious variants, only SNP rs1552726 is predicted to affect the correct splicing of gene NLRP14 [13], rs1552726 is also one of the 10 SNPs which maps reviewed genes. The left genes DPP6,Inpp5f,CCHCR1 and CBFA2T3 are also associated with many genetic diseases. Looked from the overall, we think the results are good and can validate our idea in some ways. The method can give a narrow range of suspicious pathogenic genes which deserve further studies. As whole-genome sequencing advances, more and more data can be achieved, the method can get more accurate and interesting results. The method is a simple and cheap way to find the suspicious pathogenic genes and SNPs.

